# Phosphorylation-Dependent Assembly of a 14-3-3 Mediated Signaling Complex During Red Blood Cell Invasion by *Plasmodium falciparum* Merozoites

**DOI:** 10.1101/2020.01.17.911107

**Authors:** Kunal R. More, Inderjeet Kaur, Quentin Giai Gianetto, Brandon M. Invergo, Thibault Chaze, Ravi Jain, Christéle Huon, Petra Gutenbrunner, Hendrik Weisser, Mariette Matondo, Jyoti S. Choudhary, Gordon Langsley, Shailja Singh, Chetan E. Chitnis

## Abstract

Red blood cell (RBC) invasion by *Plasmodium* merozoites requires multiple steps that are regulated by signaling pathways. Exposure of *P. falciparum* merozoites to the physiological signal of low K^+^, as found in blood plasma, leads to a rise in cytosolic Ca^2+^, which mediates microneme secretion, motility, and invasion. We have used global phosphoproteomic analysis of merozoites to identify signaling pathways that are activated during invasion. Using quantitative phosphoproteomics we found 394 protein phosphorylation site changes in merozoites subjected to different ionic environments (high K^+^/ low K^+^) out of which 143 were Ca^2+^-dependent. These included a number of signaling proteins such as catalytic and regulatory subunits of protein kinase A (PfPKAc and PfPKAr) and calcium-dependent protein kinase 1 (PfCDPK1). Proteins of the 14-3-3 family interact with phosphorylated target proteins to assemble signaling complexes. Here, using co-immunoprecipitation and gel filtration chromatography, we demonstrate that Pf14-3-3I binds phosphorylated PfPKAr and PfCDPK1 to mediate the assembly of a multi-protein complex in *P. falciparum* merozoites. A phospho-peptide, P1, based on the Ca^2+^ dependent phosphosites of PKAr, binds Pf14-3-3I and disrupts assembly of the Pf14-3-3I-mediated multi-protein complex. Disruption of the multi-protein complex with P1 inhibits microneme secretion and RBC invasion. This study thus identifies a novel signaling complex that plays a key role in merozoite invasion of RBCs. Disruption of this signaling complex could serve as a novel approach to inhibit blood stage growth of malaria parasites.

**Importance:** Invasion of red blood cells (RBCs) by *Plasmodium falciparum* merozoites is a complex process that is regulated by intricate signaling pathways. Here, we have used phosphoproteomic profiling to identify the key proteins involved in signaling events during invasion. We found changes in the phosphorylation of various merozoite proteins including multiple kinases previously implicated in the process of invasion. We also found that a phosphorylation dependent multi-protein complex including signaling kinases assembles during the process of invasion. Disruption of this multi-protein complex impairs merozoite invasion of RBCs providing a novel approach for the development of inhibitors to block the growth of blood stage malaria parasites.

## Introduction

The clinical symptoms of *Plasmodium falciparum* malaria are attributed to the blood stage of the parasite life cycle during which merozoites invade and multiply within host red blood cells (RBCs). Following the development of mature schizonts, newly formed merozoites egress and invade uninfected RBCs to initiate a new cycle of infection. The invasion of RBCs by *P. falciparum* merozoites is a complex multi-step process that is mediated by specific molecular interactions between red cell surface receptors and parasite protein ligands (1). These ligands are initially located in internal secretory organelles called micronemes and rhoptries and are released to the merozoite surface in tightly regulated steps (2).

Exposure of merozoites to a low potassium (K^+^) environment in blood plasma initiates a signaling cascade that involves second messengers like Ca^2+^ and cyclic nucleotides that activate effector molecules such as kinases and phosphatases (3–5). These effectors modulate phosphorylation of target proteins to activate merozoite motility, as well as secretion of invasion related proteins such as *P. falciparum* 175 kD erythrocyte binding antigen (PfEBA175) and apical merozoite antigen-1 (PfAMA1) from the micronemes to the merozoite surface (2). The engagement of PfEBA175 with its receptor glycophorin A triggers another signaling cascade that leads to the release of rhoptry proteins such as PfRH2b (5). The secretion of microneme and rhoptry proteins seals the engagement of the merozoite with the RBC and enables completion of the invasion process.

The signaling mechanisms that regulate processes such as apical organelle secretion and merozoite motility during host cell invasion are not fully understood. Protein phosphorylation is known to be the primary regulator of biological signaling pathways and phosphoproteome analysis can provide information about the signaling pathways that are activated in a cell in response to different stimuli (6). Protein phosphorylation/dephosphorylation acts as a molecular switch that can lead to diverse outcomes including activation or deactivation of enzymes, preparation of proteins for degradation, translocation of proteins to various cellular compartments and establishment of protein-protein interactions leading to the formation of multi-protein complexes that function in signaling pathways (7).

Phosphorylation-dependent formation of multi-protein signalling complexes plays a key role in the regulation of diverse cellular processes (8–10). For example, in human cells, phosphorylation of membrane-associated guanylate kinase-like domain-containing protein (CARMA) by protein kinase C (PKC) leads to the formation of CARMA1-Bcl10-MALT1 (CBM) complex, which activates the transcription factor NF-κB to regulate cell survival, activation and proliferation (8). A family of scaffold proteins, referred to as the 14-3-3 family, binds phosphorylated proteins to assemble signaling complexes in diverse systems (9–10). For example, in the brain, a 14-3-3ζ dimer simultaneously binds and bridges the cytoskeletal protein tau and glycogen synthase kinase, GSK3β, to stimulate tau phosphorylation, which in turn regulates microtubule dynamics (9). In the case of *Arabidopsis thaliana*, calcium-dependent phosphorylation of a basic region/leucine-zipper (bZIP) transcription factor FD leads to the formation of a florigen complex with flowering locus T protein that is mediated by 14-3-3 and regulates flowering (10). Phosphorylation analysis of *P. falciparum* schizont stages also reported the formation of a phosphorylation-dependent high-molecular-weight complex involving calcium-dependent protein kinase-1 (PfCDPK1) (11), although the precise composition of the complex was not defined.

In this study, we present a phospho-protein profile of *P. falciparum* merozoites and identify signal-dependent phosphorylation events that play important roles in the RBC invasion process. Importantly, we describe the formation of a phosphorylation-dependent, dynamic, high-molecular-weight complex involving PfCDPK1 and PfPKAr and explore the role of Pf14-3-3 in the assembly of this complex. Disruption of the Pf14-3-3-mediated protein complex with a peptide mimic inhibits RBC invasion by merozoites providing a novel strategy to block blood stage growth of malaria parasites.

## Results

### Phosphoproteome analysis of *P. falciparum* merozoites

Merozoites released from synchronized *P. falciparum* schizonts were purified (5) and processed for mass-spectrometric phosphoproteome analysis. The workflow used for phosphoproteomics and data analysis are outlined in Figure S1. Dataset S1 provides the list of phosphorylated proteins and phosphosites identified in *P. falciparum* merozoites. Comparison with the published *P. falciparum* merozoite phosphoproteome (12) identified 2786 phosphosites and 666 merozoite phosphoproteins that are unique to our study (Fig. 1a). Potential protein-protein interactions in the merozoite phosphoproteome are illustrated as MCODE clusters (Fig. S2 and Dataset S2). Enriched in MCODE cluster 1 are 132 proteins relevant to host cell invasion (Fig. 1b). The phosphorylated proteins in MCODE cluster 1 include signaling related proteins such as Protein Kinase G (PfPKG; PF3D7_1436600), Guanylate Cyclase (PfGC; PF3D7_1138400), Protein Kinase A regulatory subunit (PfPKAr; PF3D7_1223100), Protein Kinase A catalytic subunit (PfPKAc; PF3D7_0934800), Calcium Dependent Protein Kinase 1 (PfCDPK1; PF3D7_0217500) and Calcium-Dependent Protein Phosphatase Calcineurin (PfCNA; PF3D7_0802800) as well as invasion related parasite proteins such as merozoite surface protein-1 (MSP1), erythrocyte binding antigens (EBA181, EBA140) rhoptry neck proteins (RON2, RON3, RON4) and parasite proteins responsible for motility such as GAP45, GAP40, MyoA, MyoB and MTIP (Fig. 1b). The presence of both calcium and cyclic nucleotide responsive effectors in MCODE cluster 1 (Fig. 1b) indicates significant crosstalk between these second messengers at the time of invasion.

**Figure 1.**
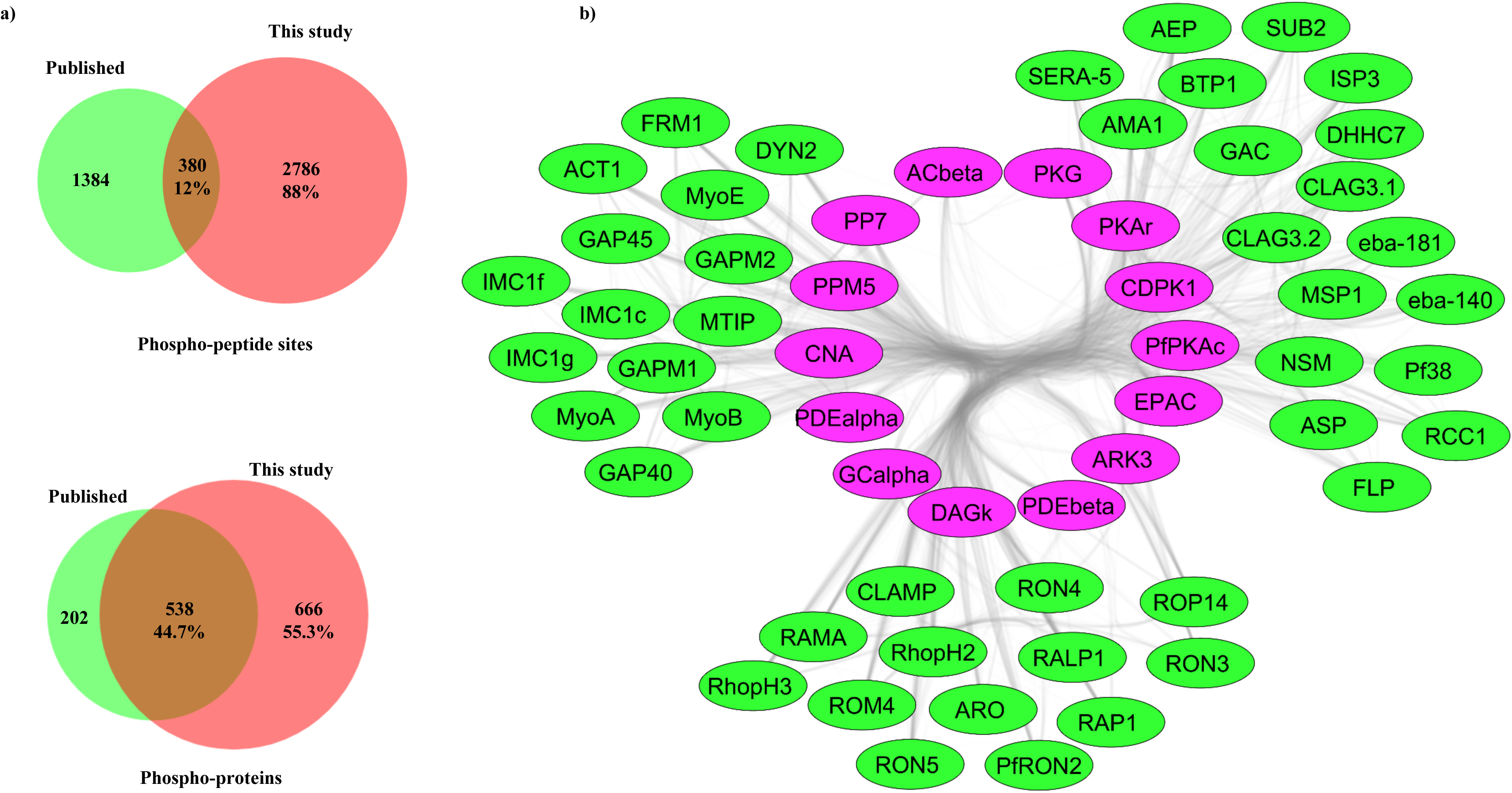
The phosphoproteome of *P. falciparum* merozoites: **a.)** Venn diagram depicting overlap in phosphorylation sites and phosphorylated proteins between the published merozoite phosphoproteome (12) and this study. **b)** Signaling and invasion related proteins from clustered subnetwork 1 from merozoite interaction network of phosphorylated proteins using MCODE clustering algorithm. Protein interaction data were downloaded from STRING for phosphoproteins found in this study and visualized in Cytoscape. MCODE clustering algorithm was used for the generation of the subnetwork of highly interacting proteins and each cluster was analyzed to identify overrepresented molecular function categories. Interaction network of proteins corresponding to molecular function category, Invasion of host cells and signaling related proteins from MCODE cluster 1, is shown here. Invasion-related proteins are in green, while signaling related proteins are in pink.

### Exposure of *P. falciparum* merozoites to an ionic environment mimicking blood plasma induces changes in protein phosphorylation

We have shown previously that merozoites respond to changes in their ionic environment, especially changes in potassium ion (K^+^) concentration (5). Exposure of merozoites to a low K^+^ environment, which is characteristic of extracellular ionic conditions in blood plasma, serves as a signal to trigger a rise in Ca^2+^ and cAMP, which activates signaling cascades (4, 5). We performed quantitative phosphoproteomics on merozoites resuspended in buffers mimicking intracellular and extracellular ionic conditions (IC buffer and EC buffer) to identify differences in protein phosphorylation. This resulted in the identification of 1499 unique phosphosites corresponding to 587 *P. falciparum* proteins (Dataset S3). Ca^2+^-dependent changes in phosphorylation were identified by studying differences in phosphorylation of merozoite proteins in IC buffer compared to either EC buffer or EC buffer + BAPTA-AM (EC-BA) (Dataset S3).

Proteins exhibiting statistically significant fold changes in phosphorylation at specific amino acid residues in merozoites in EC buffer compared to IC buffer and in EC-BA buffer compared to IC buffer were identified. Peptides from the same proteins without any phosphorylation were quantified and used to normalize for differences in concentration of proteins in merozoite samples under different conditions. Based on results of two independent biological replicates with each replicate analyzed two times by mass spectrometry, we identified 394 phosphoresidues as significantly altered when merozoites are exposed to EC buffer compared to IC buffer. Of these, phosphorylation at 143 sites is blocked by Ca^2+^ chelator, BAPTA-AM (Datasets S3 and S4). Changes in phosphorylation of some key signaling related proteins such as PfPKA-R, PfCDPK1 and Pf14-3-3I were observed (Figs. 2a, 2b). Phosphorylation of PfCDPK1 on Ser 28/34, and Ser 64 was significantly upregulated in merozoites exposed to EC buffer as compared to IC buffer (Fig. 2a, 2c). Chelation of Ca^2+^ with BAPTA-AM had no effect on these phosphorylation events (Fig. 2b, 2c). Phosphorylation on Ser 17 and Ser 217 of PfCDPK1 was found to be higher in merozoites in EC-BA buffer compared to IC buffer. However, there was no increase in phosphorylation of Ser 17 and Ser 217 in EC buffer compared with IC buffer (Fig. 2c). In contrast, phosphorylation of PfPKAr in EC buffer at Ser 113/Ser 114 was dependent on the presence of Ca^2+^ (Fig. 2d). Corresponding spectra and quantification profile for phosphorylation of PfCDPK1 on Ser 28/34 and of PfPKAr on Ser 113/Ser 114 are represented in Fig. S3. PfCDPK1 and PfPKAr are known to be involved in RBC invasion by merozoites (4, 13). We, therefore, investigated further the relevance of changes in their phosphorylation status to the process of invasion.

**Figure 2.**
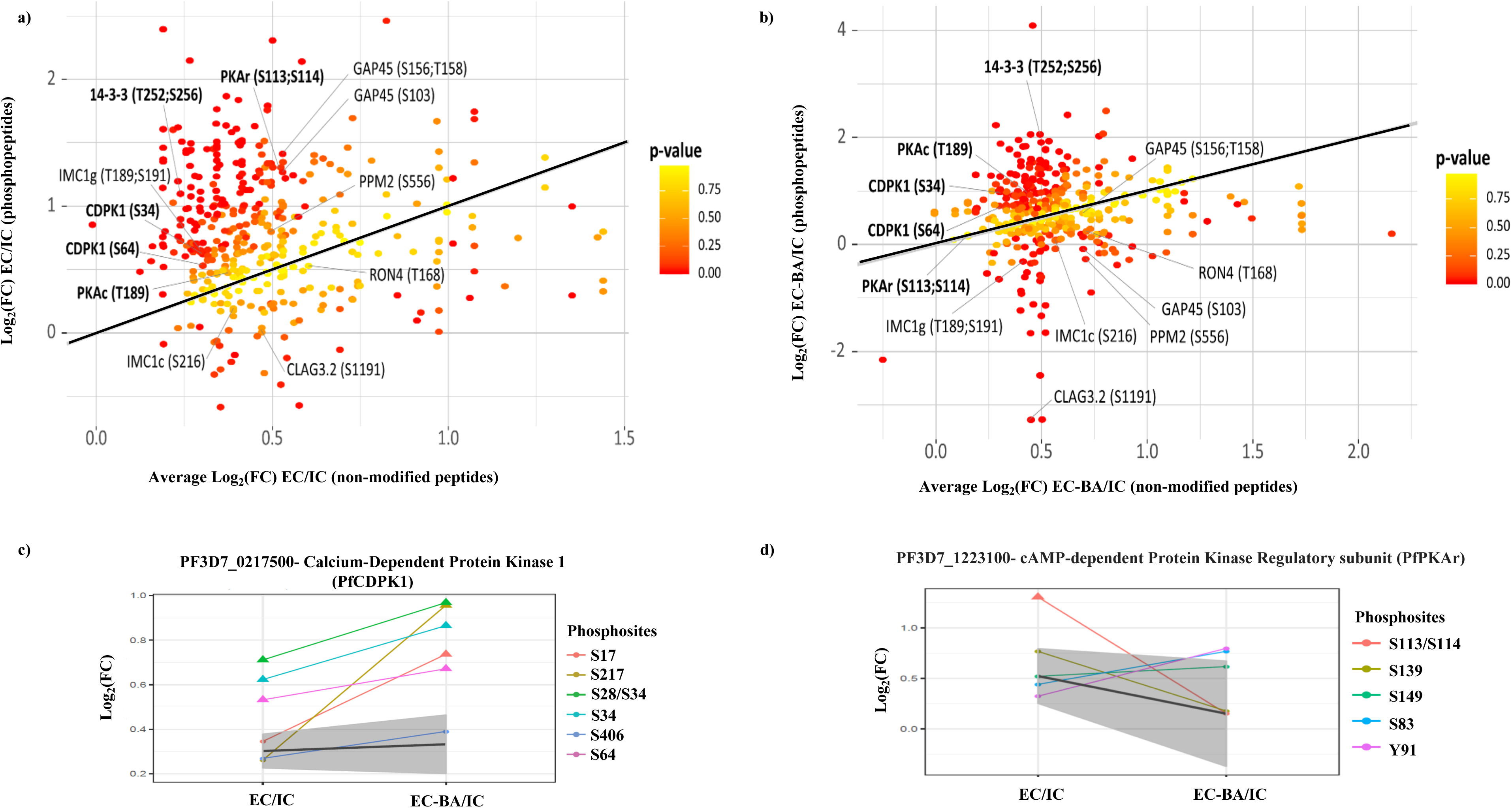
Signal-dependent changes in phosphoproteome of *P. falciparum* merozoites: **a)** Fold-changes in abundance of phosphopeptides in merozoites in EC buffer with low K^+^ compared to IC buffer with high K^+^ were plotted against fold-changes of non-phosphopeptides. **b)** Fold-changes in phosphopeptides in merozoites in EC + BAPTA-AM (EC-BA) buffer compared to IC buffer plotted against fold-changes of non-phosphopeptides. p-values for fold change in phosphorylation (colour coded) were calculated, as compared to the changes in non-phosphorylated peptides for respective proteins. Key proteins and their phosphosites with significant alterations are shown. **c.)** Fold-changes in individual phosphorylations in EC compared with IC and EC-BA compared with IC for PfCDPK1. The grey area represents fold-change in abundance of non-phosphorylated peptides from the corresponding proteins. Phosphosites with significant changes in abundance lie outside the grey area and are denoted with triangles. Phosphorylation of Ser 28/34 and Ser 64 of PfCDPK1 was significantly higher in EC and EC-BA compared to IC. **d.)** Fold-changes in individual phosphorylations in EC compared with IC and EC-BA compared with IC for PfPKAr. The grey area represents fold-change in abundance of non-phosphorylated peptides from the corresponding proteins. Phosphosites with significant changes in abundance lie outside the grey area and are denoted with triangles. Phosphorylation of PfPKAr on Ser113/Ser114 was significantly higher in EC buffer, but not EC-BA buffer, compared to IC buffer.

The changes in phosphorylation of key signaling proteins, PfCDPK1, PfPKAr and Pf14-3-3I, in EC buffer compared to IC buffer were also confirmed using anti-phosphoserine antibodies. Lysates of *P. falciparum* merozoites in IC and EC buffers were used for IP with anti-PfCDPK1 and anti-PfPKAr sera. The IPs were separated by SDS-PAGE and PfCDPK1 and PfPKAr were detected by western blotting. The blots were also probed with anti-phosphoserine antibodies to determine levels of serine phosphorylation in these proteins (Fig. S4a). Western blotting with anti-phosphoserine antibodies confirmed that the levels of phosphorylated serines in PfCDPK1 and PfPKAr were higher in EC buffer compared to IC buffer (Fig. S4a). Moreover, each IP sample showed multiple proteins with increased serine phosphorylation suggesting that these phosphorylated proteins may interact with each other to form a multi-protein complex.

### Formation of a multi-protein complex involving PfPKAr, PfCDPK1and Pf14-3-3I in *P. falciparum* merozoites

To investigate the interactions of signaling proteins, PfPKAr and PfCDPK1, in merozoites, we immunoprecipitated PfPKAr and PfCDPK1 from merozoite lysates using specific polyclonal sera and identified interacting proteins in the IPs by mass spectrometry. The presence of PfPKAr in IPs with anti-PfCDPK1 sera was confirmed by detection of multiple PfPKAr peptides with greater than 50% sequence coverage (Table 1). Similarly, presence of PfCDPK1 is confirmed in IPs performed with anti-PfKAr sera. Multiple PfCDPK1 peptides, with greater than 50% sequence coverage, are detected in IPs performed with anti-PfPKAr sera (Table 1). In addition, the scaffold protein, Pf14-3-3I, is also detected in IPs performed with both anti-PfCDPK1 and anti-PfPKAr sera with multiple Pf14-3-3I peptides detected that provide greater than 50% sequence coverage. The presence of Pf14-3-3I in the complex is further confirmed by detection of PfCDPK1 and PfPKAr peptides with greater than 50% sequence coverage in IPs performed with anti-Pf14-3-3I sera (Table 1). These studies suggest that Pf14-3-3I, PfCDPK1 and PfPKAr interact to form a multi-protein complex in *P. falciparum* merozoites (Table 1). In addition to PfPKAr, PfCDPK1 and Pf14-3-3I, four other proteins (elongation factor 1-alpha, glyceraldehyde-3-phosphate dehydrogenase (GAPDH), phosphoethanolamine N-methyltransferase (PMT), actin-depolymerizing factor 1 (ADF1)) were detected at similar stringency levels in IPs with all three sera, (anti-PfPKAr, anti-PfCDPK1 and anti-Pf14-3-3I sera) (Dataset S5). A number of other proteins are detected at lower stringency in the IPs by mass spectrometry (Dataset S5). As a negative control for specificity, we used specific antisera to *P. falciparum* protein kinase G (PfPKG) to detect its presence in IPs with anti-PfCDPK1, anti-PfPKAr and anti-14-3-3I sera. PfPKG was not detected in IP pellets with anti-PfPKAr, ant-PfCDPK1 and anti-Pf14-3-3I sera (Fig. S4b). PfPKG was also not detected in IPs with anti-PfPKAr, ant-PfCDPK1 and anti-Pf14-3-3 sera by mass spectrometry.

**Table 1.** Identification of PfCDPK1, PfPKAr, Pf14-3-3I and PfPKAc by mass spectrometry. The complete list of proteins identified in the immunoprecipitates is reported in Supplementary Table S4.

| Protein detected in IP |  |  | IP with anti-PfPKAr |  | IP with anti-PfCDPK1 |  | IP with anti-Pf14-3-3I |  |
| --- | --- | --- | --- | --- | --- | --- | --- | --- |
| protein ID | Protein name | Max Quant Score | Unique peptides | Sequence coverage [%] | Unique peptides | Sequence coverage [%] | Unique peptides | Sequence coverage [%] |
| PF3D7_0217500 | PfCDPK1 | 323.31 | 39 | 62 | 57 | 66.6 | 43 | 66.4 |
| PF3D7_1223100 | PfPKAr | 323.31 | 36 | 69.2 | 49 | 74.6 | 34 | 71.4 |
| PF3D7_0818200 | Pf14-3-3I | 323.31 | 20 | 67.2 | 19 | 58.4 | 20 | 74 |

The IPs described above were carried out with lysates made from merozoites resuspended in RPMI 1640, which has low K^+^ levels. Next, we investigated if the interactions of Pf14-3-3I, PfPKAr and PfCDPK1 are dependent on the external ionic environment and if intracellular Ca^2+^ plays a role in these interactions. Lysates of merozoites resuspended in IC, EC and EC-BA buffers were used for IP with specific sera against PfCDPK1, PfPKAr and Pf14-3-3I. The IP elutes were probed for the presence of interacting partners. PfCDPK1 and Pf14-3-3I were detected in IPs generated with specific anti-PfPKAr sera with lysates prepared from merozoites in EC buffer (Fig. 3a). In contrast, the amounts of PfCDPK1 and Pf14-3-3I in IP elutes with anti-PfPKAr sera were significantly lower in lysates prepared from merozoites in IC and EC-BA buffers. The interactions of PfPKAr with PfCDPK1 and Pf14-3-3I are thus favored when merozoites are exposed to a low K^+^ environment (Fig. 3a). Moreover, the reduced signal in IPs in case of merozoites in EC-BA buffer indicates that this interaction requires Ca^2+^. However, the interaction between Pf14-3-3I and PfCDPK1 is not dependent on presence of Ca^2+^ (Fig. 3b and 3c). Collectively, these observations suggest that PfPKAr, PfCDPK1, and Pf14-3-3I form a multi-protein complex when merozoites are exposed to a low K^+^ environment typical of blood plasma. The interaction of PfPKAr with the multi-protein complex is dependent on the presence of intracellular Ca^2+^, whereas the interaction of PfCDPK1 is independent of Ca^2+^.

**Figure 3.**
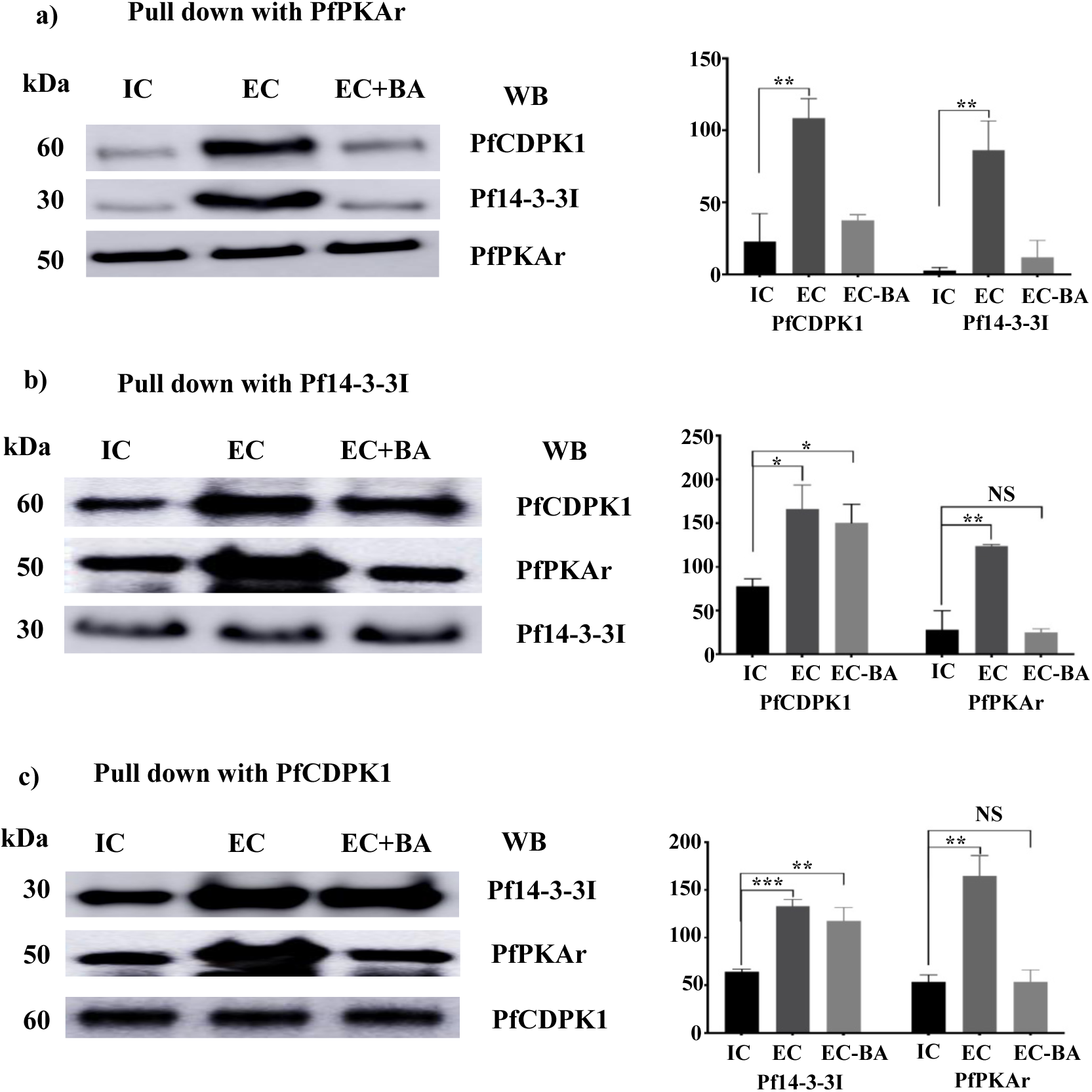
Calcium-dependent interaction of PfPKAr with PfCDPK1 and Pf14-3-3I leads to formation of a multi-protein complex in *P. falciparum* merozoites: **a)** PfPKAr was immunoprecipitated (IP) from merozoites in IC buffer mimicking intracellular ionic conditions with high K^+^ (IC), EC buffer mimicking extracellular ionic conditions with low K^+^, or EC buffer with intracellular Ca^2+^ chelator BAPTA-AM (EC-BA). The presence of PfCDPK1, Pf14-3-3I and PfPKAr in IPs was confirmed by western blotting. Graphs show the average intensity normalized with the protein immunoprecipitated for each condition from three independent experiments. A representative western blotting image from one of three independent experiments is shown. **b)** and **c)** show representative western blotting images for IPs with specific anti-Pf14-3-3I and anti-PfCDPK1 sera, respectively; with quantification of interaction partners from 3 independent experiments shown in bar graphs. Mean ± SEM are shown with n=3, * indicates P < 0.05, ** indicates P < 0.005, *** indicates P < 0.0005 by t-test. NS, non-significant, P > 0.05.

Size exclusion chromatography was also used to detect the presence of the multi-protein complex including PfPKAr, PfCDPK1 and Pf14-3-3I in merozoite lysates. When lysates were prepared from merozoites treated with IC buffer, PfPKAr, PfCDPK1 and Pf14-3-3I primarily migrated at positions reflecting their monomeric or dimeric sizes (Fig. 4). Some Pf14-3-3 and PfPKAr proteins were found in the higher molecular weight fractions in IC buffer, as 14-3-3 can exist in the form of homodimer (14) and PfPKAr interacts with PfPKAc (3). In contrast, when lysates were prepared from merozoites in EC buffer, PfPKAr, PfCDPK1 and Pf14-3-3I were primarily present in a high-molecular-weight complex migrating between 150 to 250 kDa (Fig. 4). Assembly of the PfPKAr, PfCDPK1, and Pf14-3-3I complex in merozoites is thus dynamic in nature and assembles in merozoites exposed to a low K^+^ ionic environment (Fig. 4).

**Figure 4.**
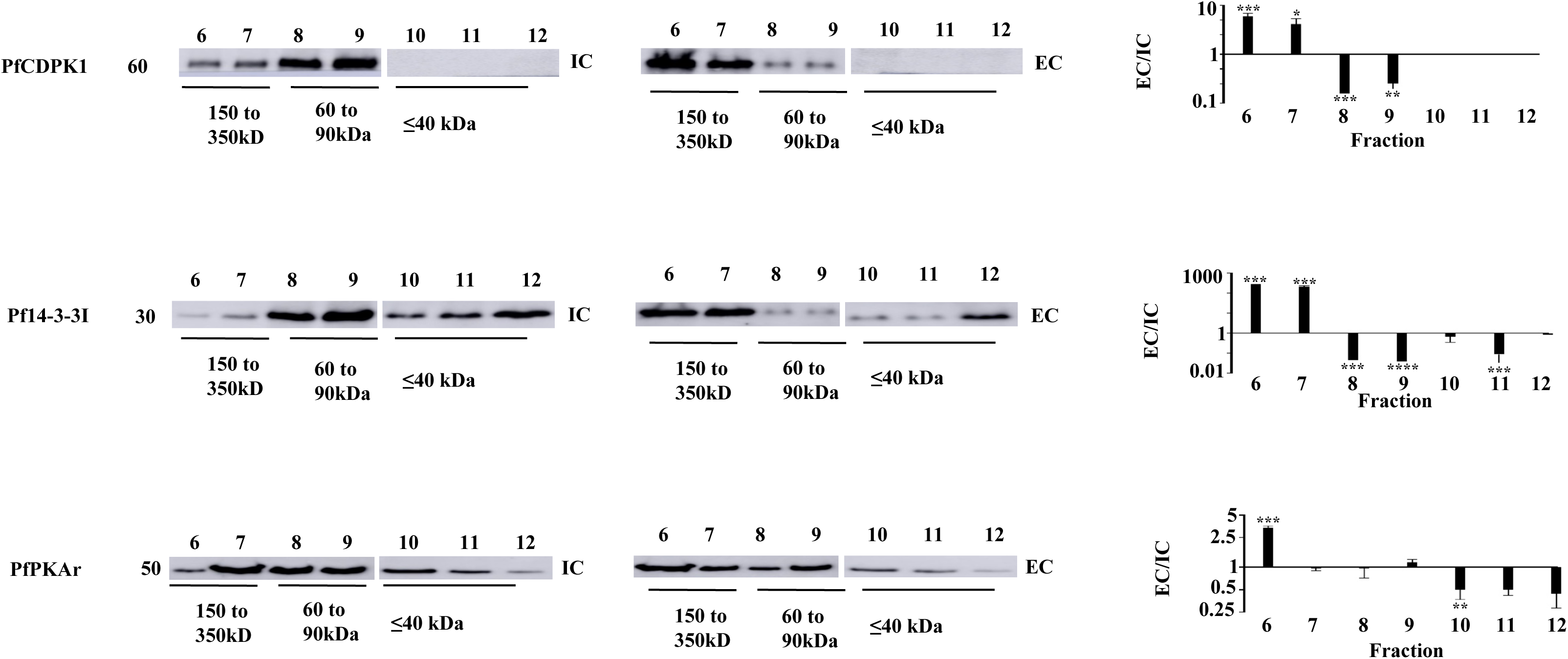
Analysis of assembly of multi-protein complexes with PfPKAr, PfCDPK1 and Pf14-3-3 by gel filtration chromatography. Lysates were prepared from merozoites in IC and EC buffers and fractionated using a Superdex 200 gel filtration column. Fractions were probed for the presence of PfCDPK1, Pf14-3-3I, and PfPKAr by western blotting. Representative blots are shown for one out of three independent experiments. Intensity for each lane (representing each fraction) was measured using ImageJ software and the ratio of EC/IC was calculated. The graph represents the average of the EC/IC ratio for each protein from three independent experiments. Mean ± SEM are shown for 3 independent experiments (n=3). * indicates P < 0.05, ** indicates P< 0.005, *** indicates P < 0.0005 by t-test.

### Recombinant Pf14-3-3I binds specifically to a phosphopeptide based on PfPKAr

Phosphorylation of PfPKAr at Ser 113 and Ser 114 following exposure of merozoites to EC buffer is dynamic and depends on intracellular Ca^2+^ levels (Fig. 2). The phosphorylation of PfPKAr and the interaction between PfPKAr and Pf14-3-3I are both dependent on the presence of Ca^2+^ (Figs. 2 and 3). As 14-3-3 family proteins are phospho-recognition scaffold proteins that participate in the formation of multi-protein complexes (9, 14), we hypothesized that interaction between Pf14-3-3I and PfPKAr requires Ca^2+^-dependent phosphorylation of PfPKAr at Ser 113 and Ser 114. To test this hypothesis, we synthesized three peptides including phosphopeptide P1 spanning the sequence of phosphorylated Ser 113 and Ser 114 (NDDG**p**S**p**SDG; P1), a non-phosphorylated peptide (NDDGSSDG; P2) and a scrambled phosphorylated P1 peptide with a random distribution of the phospho-Ser residues (pSDNGpSGDD; P3). These peptides were immobilized on agarose beads and incubated with recombinant GST-tagged Pf14-3-3I protein. There was significant binding of Pf14-3-3I-GST to beads coated with synthetic peptide P1, but none or only marginal binding to beads coated with synthetic peptides P2 and P3 (Fig. 5a). Known 14-3-3 binding peptides AA (ARSHpSYPA) and RA (RLYHpSLPA) based on canonical 14-3-3 binding motifs (15) also showed binding to Pf14-3-3I-GST similar to peptide P1 in control experiments (Fig. 5a).

**Figure 5.**
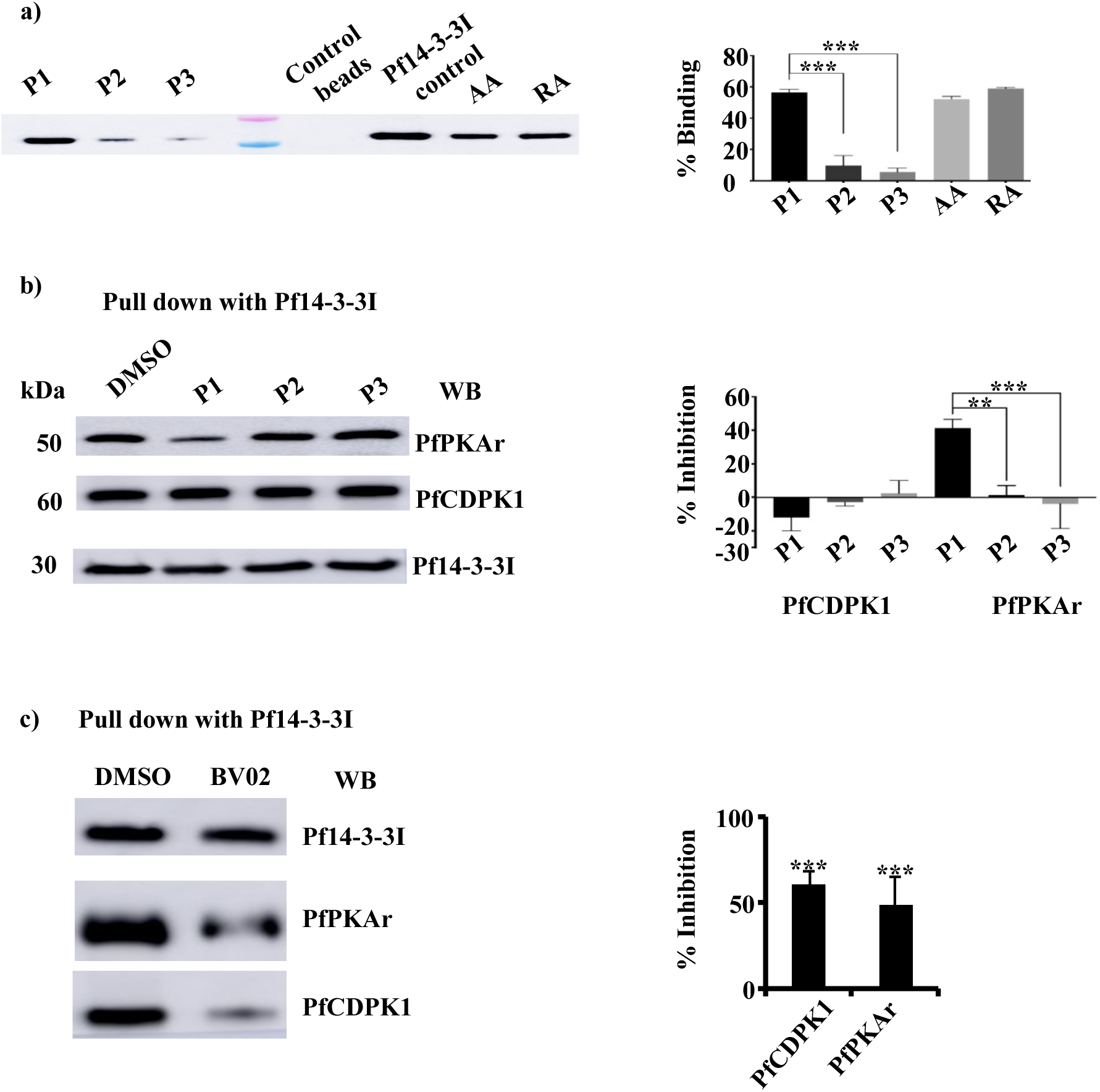
Inhibition of interaction between PfPKAr and Pf14-3-3I by a PfPKAr-derived phosphopeptide, P1, and small-molecule inhibitor, BV02, of Pf14-3-3I. **a)** Binding of Pf14-3-3I with phosphopeptide P1 from PfPKAr. Phosphopeptide P1 (NDDGpSpSDG) based on amino acid sequence of calcium-dependent phosphosites on PfPKAr encompassing S113 and S114, non-phosphorylated control peptide P2 (NDDGSSDG) and peptide P3 (pSDNGpSGDD), a control phosphopeptide based on scrambled P1 sequence, were tested for binding to Pf14-3-3I. Peptides P1, P2 and P3 were immobilized on agarose beads and allowed to interact with recombinant Pf14-3-3I. Bound recombinant Pf14-3-3I was detected in elutes by western blotting. Phosphopeptides AA and RA, which are known to bind 14-3-3 were used as positive controls. Control beads with no immobilized peptides were used as negative control. Plots show the average percentage binding calculated for binding of recombinant Pf14-3-3I to each peptide from 3 independent experiments. A representative western blotting image from one out of three independent experiments is shown. **b)** Peptide P1 inhibits binding of Pf14-3-3 to PfPKAr in merozoites. Merozoites were treated with 100 μM P1, P2, P3, or RPMI. Merozoites were lysed and lysates were used for immunoprecipitation with anti-Pf14-3-3I sera. Presence of PfCDPK1 and PfPKAr in the immunoprecipitates was investigated by western blotting. Percent inhibition of binding of PfCDPK1 and PfPKAr with Pf14-3-3I was calculated. The plot shows average percent inhibition of binding for three independent experiments. P1 decreases binding of PfPKAr to Pf14-3-3I, but has no effect on PfCDPK1 binding to Pf14-3-3I. **c)** BV02, a small molecule inhibitor of 14-3-3 interactions with phosphopeptides, inhibits binding of Pf14-3-3I to PfCDPK1 and PfPKAr in merozoites. Merozoites were treated with 2 μM BV02 or DMSO and lysed, lysates were used for immunoprecipitation with specific anti-Pf14-3-3I serum. Presence of PfCDPK1 and PfPKAr in the immunoprecipitates was confirmed by western blotting. The plot shows the mean percent inhibition of binding (± SEM) for three independent experiments (n=3). * indicates P < 0.05, ** indicates P < 0.005, *** indicates P < 0.0005, by t-test. NS, not significant (P > 0.05).

### Phosphopeptide from PfPKAr specifically inhibits interaction between Pf14-3-3I and PfPKAr in merozoites

Given that phosphopeptide P1 can bind to Pf14-3-3I *in vitro,* we next tested if P1 can inhibit binding of Pf14-3-3I to PfPKAr to inhibit multi-protein complex formation in merozoites. We first confirmed that peptide P1 can enter *P. falciparum* merozoites. Peptide P1 tagged with fluorophore fluorescein isothiocynate (FITC) was incubated with merozoites and tested for uptake by detecting the internalized peptide by fluorimetry (Fig. S5). Uptake of P1-FITC was observed at concentrations above 25 µM (Fig. S5). Merozoites were incubated with peptides P1, P2 and P3 at 100 µM. Subsequently, merozoite lysates were used for IP with anti-Pf14-3-3I antisera. PfPKAr and PfCDPK1 were detected in the IPs by western blotting. Phosphorylated peptide P1 inhibited the interaction between Pf14-3-3I and PfPKAr, while peptides P2 and P3 had no effect (Fig 5b). Interestingly, none of the peptides (P1, P2, and P3) had any effect on the interaction between Pf14-3-3I and PfCDPK1. The small-molecule inhibitor BV02 that blocks binding of mammalian 14-3-3 to its phosphorylated target proteins (15, 16), inhibited binding of both PfPKAr and PfCDPK1 with Pf14-3-3I (Fig. 5c).

### Blocking Pf14-3-3I interactions inhibits merozoite invasion of RBCs and microneme secretion

We observed above that a low K^+^ environment triggered Pf14-3-3I mediated formation of a high-molecular-weight multi-protein complex composed of PfCDPK1, PfPKAr, and PKAc, all of which are involved in RBC invasion (4, 5, 17). Next, we investigated if disruption of Pf14-3-3I-mediated binding of PfPKAr and PfCDPK1 can inhibit RBC invasion. *P. falciparum* merozoites isolated in low K^+^ buffer were treated with increasing concentrations of peptides P1, P2 and P3 (10 µM, 50 µM or 100 µM) and BV02 (0.5 µM, 1 µM, 1.5 µM or 2 µM). Treated merozoites were then incubated with RBCs in complete RPMI medium to allow invasion. Newly invaded ring-stage parasites were scored by flow cytometry. Treatment of merozoites with peptide P1 and BV02 reduced the efficiency of erythrocyte invasion in a dose-dependent manner (Fig. 6a). Control peptides, P2 (without phosphorylation) and P3 (scrambled phosphopeptide) had no inhibitory effect on invasion.

**Figure 6.**
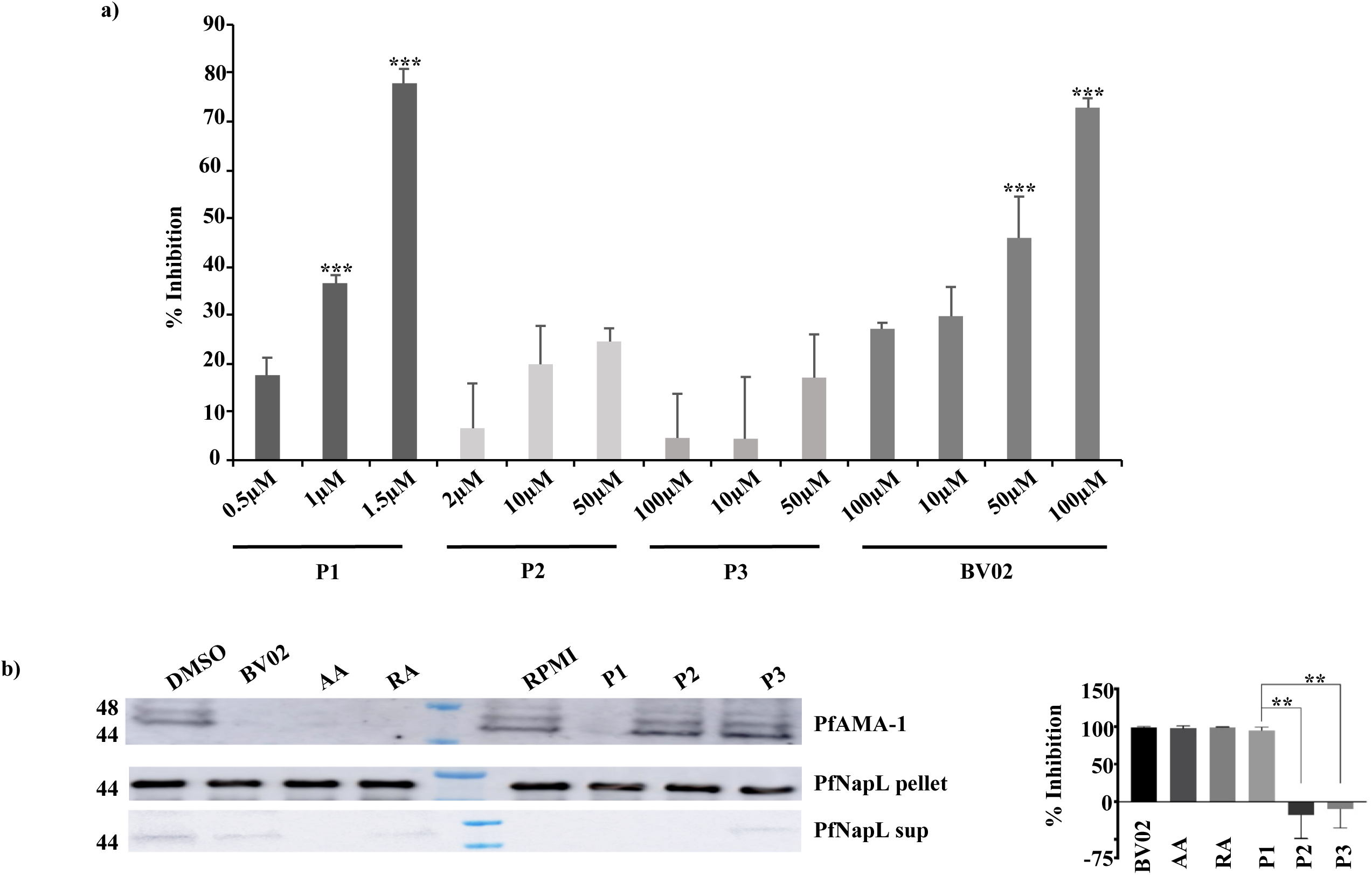
Inhibition of RBC invasion and microneme secretion by both PfPKAr-based phosphopeptide and Pf14-3-3I inhibitor. **a)** Peptide P1 and BV02 both block RBC invasion by merozoites. *P. falciparum* merozoites were isolated and allowed to invade erythrocytes in the presence of increasing concentrations of peptides P1, P2 amd P3 (10, 50 and 100 µM) and increasing concentrations of BV02 (0.5, 1, 1.5 and 2 µM). Newly invaded trophozoites were stained with SYBR green and scored by flow cytometry. Merozoites were allowed to invade erythrocytes in the absence of inhibitors using respective solvents as control. Percent invasion inhibition rates (Mean ± SEM with n=3) in presence of inhibitors are shown. ** indicates P < 0.005, *** indicates P < 0.005, t-test. **b)** Phosphopeptide P1 (NDDGpSpSDG) derived from the amino sequence of 2 calcium-dependent phosphosites on PfPKAr encompassing S113 and S114, non-phosphorylated control peptide P2 (NDDGSSDG) and peptide P3 (pSDNGpSGDD), a control phosphopeptide based on scrambled P1 sequence, AA and RA, 2 phosphopeptides based on 14-3-3 substrate binding sites, and BV02, a small molecule 14-3-3 binding inhibitor were tested for inhibition of PfAMA1 secretion by merozoites. Secretion of PfAMA1 was significantly reduced upon treatment of merozoites with phosphopeptides P1, AA and RA, and by BV02. Cytoplasmic protein PfNAPL was detected in the supernatant and used as a control for merozoite lysis. PfNAPL was detected in merozoite pellet and used as a control for normalization. Western blotting image for one out of three independent experiments is shown. The plot shows the average percent inhibition of binding for three independent experiments. Mean ± SEM are shown (n=3), ** indicates P < 0.005 by t-test.

**Figure 7.**
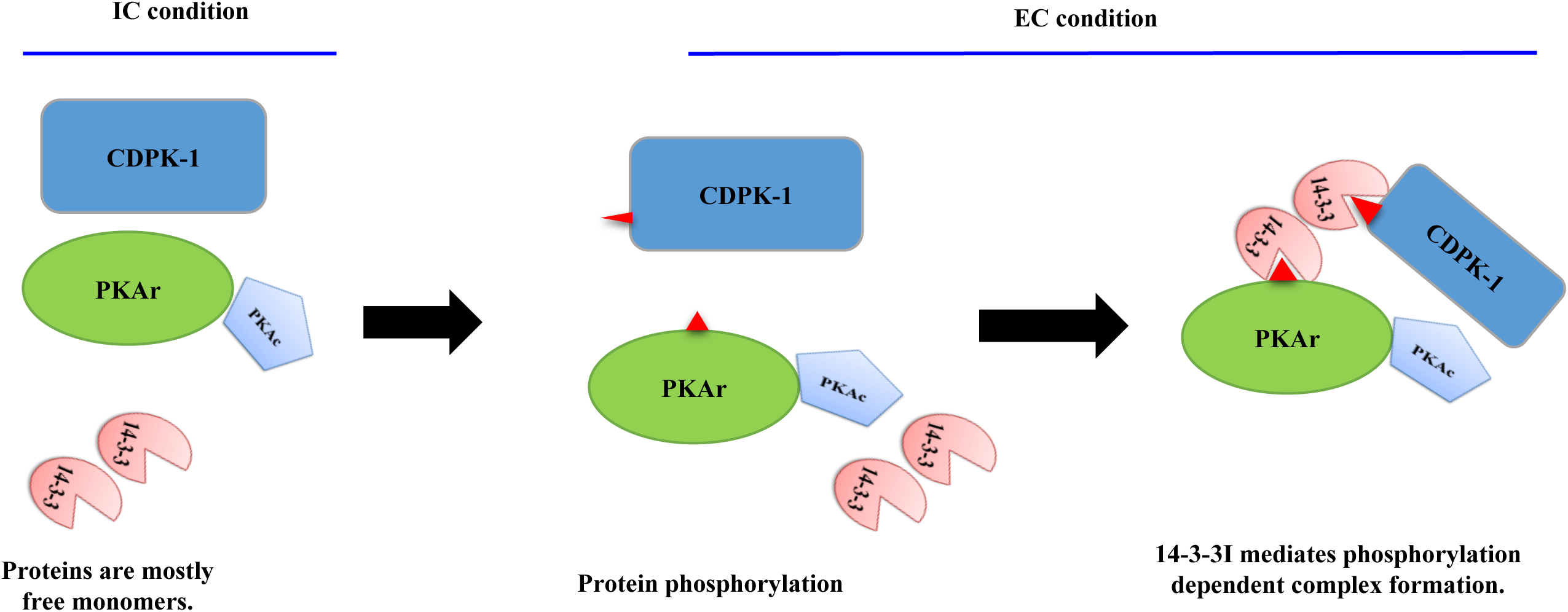
Model for assembly of a signaling complex during RBC invasion by *P. falciparum* merozoites. Exposure of merozoites to a low K^+^ ionic environment as found in blood plasma triggers a signaling cascade resulting in the phosphorylation of PfCDPK1 and PfPKAr. The scaffold protein Pf14-3-3I binds phosphorylated PfPKAr and PfCDPK1 leading to the formation of a high-molecular-weight multi-protein complex composed of PfCDPK1, PfPKAr, PfPKAc and Pf14-3-3. Formation of this signalling complex plays a regulatory role in secretion of microneme proteins and merozoite invasion of RBCs.

The secretion of parasite invasion ligands from micronemes is a critical step in the invasion process (2). Given that PfCDPK1 and PfPKAr have both been implicated in microneme secretion (4, 5), we examined if disruption of the high-molecular-weight multi-protein complex composed of PfCDPK1, PfPKAr, and PKAc, can disrupt microneme secretion. Treatment of merozoites with peptide P1 inhibited secretion of microneme protein PfAMA1, whereas control peptides P2 and P3 had no effect (Fig. 6b). BV02 and peptides AA and RA also inhibit PfAMA1 discharge (Fig. 6b). Formation of the Pf14-3-3I-mediated multi-protein complex, which includes PfCDPK1 and PfPKAr, thus appears to be important for regulation of microneme secretion, a key step in the invasion process.

## Discussion

Though there are many studies describing the intraerythrocytic phosphoproteomic profiles of *P. falciparum* blood stages, only one previous study described the phosphoproteome of free merozoites (12). However, it did not address dynamic changes in the merozoite phosphoproteome during the key process of RBC invasion. Here, we have investigated changes in the phosphoproteome of free merozoites following exposure to the environmental signal of low K^+^ as found in blood plasma during the process of invasion following egress from schizonts.

The study identified signal-dependent changes in phosphorylation of merozoite proteins shedding light on molecular events that govern RBC invasion. Gene Ontology and protein-protein interaction analyses predicted that a significant number of the phosphorylated proteins identified in merozoites have potential roles in signaling and invasion related processes. Analysis of protein interaction data indicated that various proteins from these categories cluster together (Fig. 1b). Of these, the most notable invasion related proteins are inner membrane complex (IMC) proteins like PfIMC1c and PfIMC1g that are associated with merozoite motility (1). These proteins connect to myosin filaments that form the conserved molecular machinery for merozoite motility that is necessary for invasion. Various myosin molecules, PfMyoA, PfMyoB and PfMyoE, as well as glideosome-associated proteins, PfGAP40 and PfGAP45, were phosphorylated. It has been proposed recently that regulated phosphorylation of S19 in PfMyoA may enhance force generation during parasite invasion (18).

At the time of merozoite egress and re-invasion, signaling cascades are initiated through the generation of second messengers including Ca^2+^, cAMP, and cGMP (2, 3). We found that various kinases and phosphatases in MCODE cluster 1, which are regulated by these secondary messengers and are associated with the process of secretion and invasion, are phosphorylated (Fig 1b). For example, calcium-dependent protein kinase PfCDPK1, PfPKAc, PfPKAr, PfPKG and the calcium dependent phosphatase, PfCNA (3,11,12,17,19,20), were phosphorylated. Our previous studies have demonstrated that exposure of merozoites to a low K^+^ environment in blood plasma following egress results in a rise in cytosolic Ca^2+^, which initiates the release of invasion-related microneme proteins (5). We investigated changes in protein phosphorylation following the transfer of free merozoites from a buffer with high K^+^ (IC buffer) to one with low K^+^ (EC buffer). The role of intracellular Ca^2+^ in phosphorylation was also investigated by transferring merozoites to EC buffer with the intracellular Ca^2+^ chelator, BAPTA-AM (EC-BA). Phosphorylation changes occurred at 394 sites located on 314 peptides, with 76 peptides having dual phosphorylations when merozoites were transferred from IC to EC buffer. Out of these 143 phosphorylation events were found to be Ca^2+^-dependent and were located on 119 peptides, 24 of which displayed dual phosphorylations. The employment of IMAC-based enrichment identified many dual phosphorylation events, which were not previously reported. Such dual phosphorylations can influence the activity of kinases. For example, dual phosphorylation of extracellular signal-regulated kinase 2 (ERK2) increases its activity by 10- to 100-fold (21). We observed many calcium-dependent dual phosphorylations on proteins with diverse functions in the life cycle of the parasite. These included proteins known to be involved in organelle secretion and invasion-related processes. For example, dual phosphorylation of inner membrane complex (IMC) protein, PfIMC1g, at Thr189/Ser191 and Tyr272/Ser274, and the glideosome−associated protein 45 (PfGAP45) at Ser156/Thr158 was observed. Both these proteins are known to be phosphorylated by a calcium-dependent kinase, PfCDPK1 (17). We also observed Ca^2+^-dependent dual phosphorylation of PfPKAr at Ser113/Ser114, but by contrast dual phosphorylation at Ser28/Ser34 of PfCDPK1 was not Ca^2+^-dependent.

Interactions between PfPKAr and PfCDPK1 and between PfPKAr and Pf14-3-3I have been observed previously (17, 22). Here, we demonstrated that these interactions are phosphorylation-dependent and dynamic (Fig. 3). Moreover, they lead to the formation of a high-molecular-weight (150-250 kDa) multi-protein signaling complex that assembles in response to a change in the environmental ionic composition to which egressed merozoites are exposed (Fig 4d). PfCDPK1 was previously shown to be present in merozoites in a high-molecular-weight complex, but the composition and dynamic nature of the complex was not described (11). A related study has shown by immunofluorescence microscopy that Pf14-3-3I colocalizes with PfCDPK1 at the periphery of merozoites (23). Our mass spectrometric analyses demonstrated that the complex contains PfCDPK1, Pf14-3-3I, PfPKAr, and PfPKAc, but interestingly, it failed to detect PfPKG that has also been implicated in apical organelle secretion at the time of merozoite invasion (24). A previous study on protein signaling complexes also did not detect PfPKG in the high-molecular-weight complex with PfCDPK1 (11).

Members of the 14-3-3 family of scaffold proteins bind target proteins in a phosphorylation-dependent manner through recognition of optimal consensus sequence motifs corresponding to mode-I (RXXpS/pT), mode-II (RXXXpS/pT) or mode-III (RXXpS/pTX1-2 C’), thus regulating a wide variety of cellular processes (25–28). Disruption of the interactions mediated by 14-3-3 proteins results in ablation of key cellular processes (17). Here, we show that peptides based on the dual phosphorylation of PfPKAr at Ser 113 and Ser 114 (peptide P1), as well as inhibitory peptides, AA and RA, that are based on consensus sequences in 14-3-3 binding proteins from mammalian cells, and 14-3-3 based inhibitory small molecule, BV02, all inhibited the formation of the multi-protein complex (Fig. 5b, c). The inhibitory peptides and BV02 also blocked secretion of microneme protein PfAMA1 and RBC invasion, demonstrating that assembly of this signaling complex plays a critical role in these processes (Fig. 6). A related study has confirmed the phosphorylation dependent interaction of recombinant Pf14-3-3I and PfCDPK1 using ELISA plate based binding assays as well as surface plasmon resonance (SPR) and isothermal calorimetry (ITC) (23). Moreover, the study shows that peptides AA and RA inhibit PfCDPK1 binding with Pf14-3-3I and block blood stage parasite growth (23). Here, we demonstrate that phosphopeptides, P1 and P2, that are based on PfPKAr sequences that interact with Pf14-3-3I, inhibit interaction of PfPKAr with Pf14-3-3I and block miconeme secretion and RBC invasion by *P. falciparum* merozoites.

14-3-3 homologs in mammalian cells are known to serve as the central hub for signaling networks that regulate cell proliferation, adhesion, survival, and apoptosis (15). Given its central role in cell growth, 14-3-3 is also implicated in the development of cancer, and small molecule inhibitors that target its scaffold function are being developed for cancer therapy (29). In this study, we have shown that targeting the assembly of the multi-protein complex in merozoites mediated by Pf14-3-3I provides a novel strategy to inhibit the blood-stage growth of malaria parasites.

## Materials and Methods

### *P. falciparum* merozoite isolation

*P. falciparum* 3D7 blood stages were cultured *in vitro* and merozoites were isolated as previously described (5, 30). Mature synchronized schizonts were transferred to IC buffer (142 mM KCl, 5 mM NaCl, 2 mM EGTA, 1 mM MgCl_2_, 5.6 mM glucose, 25 mM Hepes, pH 7.2). Released merozoites were collected by centrifugation as described previously (5) and resuspended in IC buffer, EC buffer (5mM KCl, 142 mM NaCl, 1 mM CaCl_2_, 1 mM MgCl_2_, 5.6 mM glucose and 25 mM Hepes, pH 7.2) or EC-BA buffer (EC buffer supplemented with 50 mM BAPTA-AM (Calbiochem)) at 37^0^C for 15 mins with or without inhibitors as required. Merozoites pellets were prepared by centrifugation at 3300g for 5 min and stored at −80°C for further analysis.

### Protein isolation, desalting, and digestion

Isolated merozoites were lysed by incubation with urea lysis buffer on ice for 15 min followed by sonication for 3 x 30 seconds on ice. Protein concentration was quantified by Pierce™ BCA Protein Assay Kit as per the supplier’s protocol. 6-7 mg of total protein was used for each biological replicate for the label-free phosphoproteomics experiment. For the quantitative experiment, 100 μg of total protein isolated from IC, EC, and EC-BA treated merozoites was used for labeling as described below. The isolated proteins were reduced, alkylated and digested with trypsin gold (Promega) with 1:200 enzyme to substrate ratio, at 37°C overnight. Tryptic digested peptides from the label-free experiment were desalted with reverse-phase tC18 SepPak solid-phase extraction cartridge 500mg (Waters) as described previously (31).

### Ion-exchange fractionation of *P. falciparum* merozoite lysates

Fractionation of desalted peptides was carried out with strong cation exchange (SCX) chromatography on a polySULFOETHYL-A column as described before (31). 12-15 fractions of 4ml were collected, lyophilized till the volume was reduced to 30% and desalted as described above.

### Tandem Mass Tag (TMT) labeling and fractionation using hydrophilic interaction liquid chromatography (HILIC)

Proteins isolated from merozoites in IC, EC, and EC-BA buffers were digested with trypsin and peptides were labeled separately with TMT tags with mass 128, 129 and 130 respectively as per the manufacturer’s instructions. Labeled peptide samples were combined and fractionated by HILIC using method described previously (32). 12-15 fractions (0.5 ml) were collected and lyophilized for further use.

### Immobilized metal affinity chromatography (IMAC) and TiO_2_/ZrO_2_ phosphopeptide enrichment and desalting

Combined IMAC based phosphoproteomic enrichment and desalting was carried out as described previously (31). Peptide fractions were incubated on a rotating platform with IMAC beads (PHOS-Select iron affinity gel (Sigma)) for 1hr at RT. During this time StageTips (33) were prepared using Empore 3M C18 material (Fisher Scientific). After incubation, IMAC beads were added on top of the StageTips and the flow-through was collected and concentrated on a speedvac. Phosphopeptides were eluted from IMAC resin onto C18 loaded tips and desalted. Phosphopeptides were eluted from C18 StageTips, lyophilized and stored at −80°C. Flow through from IMAC was used further for phosphopeptide enrichment using TiO_2_/ZrO_2_ NuTip (Glygen) as per manufacturer’s protocol. Phosphopeptides were eluted, concentrated by speedvac centrifugation and stored at −20°C till further analysis.

### Liquid chromatography-tandem mass spectrometry

LC-MS/MS was carried out using a Nano LC-1000 HPLC nanoflow system (Thermofisher Scientific) and hybrid Orbitrap Velos Pro mass spectrometer (Thermofisher Scientific). Peptides were separated by a 120 min gradient using Acclaim® PepMap100 C18 column and eluted onto the mass spectrometer. Data acquisition was performed in a data-dependent mode to automatically switch between MS, MS_2_. Full-scan MS spectra of intact peptides (m/z 350–1000) were acquired in the Orbitrap with a resolution of 60,000. Top 20 precursors were sequentially isolated and fragmented in the high-energy collisional dissociation (HCD) cell. Dynamic exclusion was 50 s and a minimum 500 counts for Mz and 200 counts for TMT sets were required for MS_2_ selection.

### Data analysis for TMT and label free phosphoproteomic data

All raw files were searched against a *P. falciparum* database using an OpenMS pipeline (34) containing the two search engines Mascot and MSGF+, followed by Percolator post-processing and phosphorylation analysis using PhosphoScoring, an implementation of the Ascore algorithm (35). Search parameters were: carbamidomethylation of cysteines was set as a fixed modification, oxidation of methionine, protein N-terminal acetylation, and STY phosphorylation were set as variable modifications. The mass tolerances in MS and MS/MS were set to 20 ppm and 0.5 Da respectively. A false discovery rate of 1% was set up for both protein and peptide levels. TMT experiment, TMT-6plex labeling on lysine and N-termini was searched for protein quantitation. Phosphoscore more than 11 was considered as a significant localization score.

Data was also searched using MaxQuant (version 1.5.3.8) (with the Andromeda search engine) against a *P. falciparum* database. The following search parameters were applied: carbamidomethylation of cysteines was set as a fixed modification, oxidation of methionine, protein N-terminal acetylation, and STY phosphorylation was set as variable modifications. The mass tolerances in MS and MS/MS were set to 5 ppm and 0.5 Da respectively. A false discovery rate of 1% was set up for both protein and peptide levels. TMT-6plex labeling on Lysine and N-termini was searched for protein quantitation. Phospho-localization probability of more than 75% was considered as significant localization. Quantification from MaxQuant analysis was used for quantification of changes in the phosphorylation. All phospho-spectra of interest were manually validated.

To determine whether the variation of the quantification of a phosphopeptide is due to a variation in the abundance of the protein itself, or due to a variation in the abundance of its modification, a statistical test was performed to compare the variation in abundance of each phosphopeptide to the abundance of the corresponding protein. To do this for a specific phosphopeptide, we first estimated the average *m* and standard deviation *s* of the log_2_ fold change (log_2_FC) of the non-modified peptides of the protein (where FC is equal to either 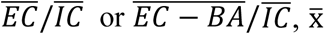 being the average intensity observed for a peptide in the condition x). Assuming that the log_2_FC of the non-modified peptides follows a Normal distribution centered on *m* and having a standard deviation of *s*, we deduce a p-value related to the test that the measured log_2_FC for the phosphopeptide is equal to *m* by 2 × *P_N(m,s)_* (log2(FC)) if log2(FC) < m and 2 × (1 − *P_N(m,s)_* (log2(FC))) if log2(FC) ≥ m, where *P_N(m,s)_* is the cumulative distribution function of *N*(*m*, *s*). Note that this p-value is computed only when we have at least 3 non-modified peptides with intensity values for a protein.

### Phosphorylated protein interaction network analysis

The merozoite phosphoproteome interaction network was constructed using STRING database and visualized in Cytoscape version 3.7.1 (36). The merozoite phospho-interactome was analyzed for highly connected nodes with the molecular complex detection clustering algorithm MCODE (37).

### Gene ontology and motif analysis

All *P. falciparum* gene ontology analyses were performed with the inbuilt result analysis tool for gene ontology on PlasmoDB database. Phosphopeptides with a width of 15 amino acids were subjected to motif analysis using MotifX (38, 39). A background of *P. falciparum* protein database was used for the analysis and occurrences threshold was set to default P-value threshold of ≤ 1e^-6^ was used to identify enriched motifs.

### Immunoprecipitation (IP), LC-MS/MS and data analysis

IP of proteins from merozoites isolated in cRPMI or resuspended in IC, EC, and EC-BA buffers or treated with specific inhibitors or peptides were performed with Pierce Co-Immunoprecipitation (Co-Ip) Kit (Pierce) as per the manufacturer’s protocol. Identity of proteins from the respective elute of IP experiments was investigated using an Orbitrap Q Exactive Plus mass spectrometer (Thermofisher Scientific) or by western blotting. Data were searched using MaxQuant as described above.

### Gel-filtration on Superdex 200

*P. falciparum* merozoites treated with IC or EC buffer were lysed and cleared by centrifugation. Proteins were fractionated with Superdex 200 column (GE Healthcare, 10×300 mm). Fractions of 1 ml were collected and analyzed by western blotting for the presence of PfPKAr, Pf14-3-3I, and PfCDPK1 in respective fractions.

### Binding of recombinant Pf14-3-3I to synthetic peptide-coated beads

Peptides (AA, RA, P1, P2, and P3) were coupled to agarose beads using Co-IP kit (Pierce) as per manufacturer’s instruction. Respective peptide coated beads were incubated with GST tagged Pf14-3-3I protein, non-coated beads were used as a control. Recombinant protein bound to the beads were eluted and elutes were tested by western blotting using anti-Pf14-3-3I mouse sera.

### Invasion assay *with P. falciparum* merozoites and flow cytometry

Merozoites isolated as described above were treated with inhibitors (BV02 at 0.5µM, 1µM, 1.5µM, 2µM) and peptides (P1, P2, P3 at 10µM, 50µM, 100µM) for 15 min at 37°C followed by incubation with RBC in presence of the inhibitor or peptide to allow invasion and growth for 24 h under standard culturing conditions. Solvents used to dissolve the inhibitors were used as control. The parasitemia was determined by flow cytometry after staining with SYBR-GreenI (Sigma) as described (40). Data were analyzed with FlowJo (Tree Star) and percent inhibition of invasion was calculated using the formula: (1-T/C) x 100; where T and C denote parasitemia in treatment and control samples respectively.

### Microneme secretion assay

*P. falciparum* merozoites isolated in cRPMI were incubated for 15 min at 37°C with BV02 (2µM) or AA (100µM) or RA (100µM) or DMSO (solvent) and with Peptides P1, P2, P3 (100µM each) or RPMI (solvent). Following incubation, merozoites and supernatants were separated by centrifugation and the presence of PfAMA1 (microneme protein), and PfNapL (cytosolic protein used as lysis control) in the supernatant, and PfNapL in the pellet (cytosolic protein used as a loading control) were detected by western blotting as described above.

### Densitometry and statistical analysis

Image J (NIH) software was used to perform densitometry of western blots. The band intensity of the loading control was used for normalization. Statistical analysis for all the plots was performed using GraphPad Prism 8.1.2 software. All experiments were analyzed using multiple *t*-test (assuming equal standard error (SE)) and P≤ 0.05 was considered significant, *= P< 0.05; **= P< 0.005; and ***= P< 0.0005. The graphs were plotted with a mean ± SEM of the population.

### Data availability

The mass spectrometry-based proteomics data have been deposited to the ProteomeXchange Consortium (http://proteomecentral.proteomexchange.org) via the PRIDE (41) partner repository with the data set identifier PXD015093.

Reviewer account details: Username; Password: MNKnPy0v

All other relevant data are available from the authors upon request.

## Acknowledgments

This work was supported by MolSigMal grant (ANR-17-CE15-0010) from Agence Nationale de la Recherche (ANR) to CEC and internal funds from the Institut Pasteur. GL acknowledges support from the Labex ParaFrap (ANR-11-LABX-0024). KRM received a Short Term Fellowship from European Molecular Biology Organization (EMBO).

## Author Contributions

KRM designed and performed all experiments, analyzed data and wrote the first draft of the manuscript; IK performed MS/MS experiments and preliminary analysis; QGG and BI performed bio-informatic and statistical analysis of phosphoproteomics data; TC performed MS/MS analysis of immunoprecipitation experiments; RJ produced reagents (antibodies to Pf14-3-3 and plasmid construct for expression of recombinant Pf14-3-3I); CH produced recombinant Pf14-3-3 for use in binding experiments and performed GPC to detect multi-protein complexes in parasite lysates; MM supervised MS/MS experiments and data analysis; HW and PG helped with analysis of phosphoproteomics data; JC supervised MS/MS data analysis; GL provided reagents (antibodies against PfPKAr and PfPKAc), helped design experiments and analyze data and edited the manuscript; SS contributed reagents (antibodies to Pf14-3-3I, plasmid construct for production of recombinant Pf14-3-3I), helped design experiments and analyzed data; CC created the project, raised funds, designed experiments, analyzed data and edited the manuscript. All authors reviewed and commented on the manuscript.

